# Key histone chaperones have distinct roles in replisome progression and genomic stability

**DOI:** 10.1101/2020.02.25.964221

**Authors:** Ioannis Tsirkas, Daniel Dovrat, Yang Lei, Angeliki Kalyva, Diana Lotysh, Qing Li, Amir Aharoni

**Affiliations:** Department of Life Sciences and the National Institute for Biotechnology in the Negev, Ben-Gurion University of the Negev, Be’er Sheva 84105, Israel; State Key Laboratory of Protein and Plant Gene Research, School of Life Sciences and Peking-Tsinghua Center for Life Sciences, Peking University, Beijing 100871, China

## Abstract

Replication-coupled (RC) nucleosome assembly is an essential process in eukaryotic cells in order to maintain chromatin structure during DNA replication. The deposition of newly synthesized H3/H4 histones during DNA replication is facilitated by specialized histone chaperones. Although the contribution of these histone chaperones to genomic stability has been thoroughly investigated, their effect on replisome progression is much less understood. By exploiting a time-lapse microscopy system for monitoring DNA replication in individual live cells, we examined how mutations in key histone chaperones including *CAC1*, *RTT106*, *RTT109* and *ASF1*, affect replication fork progression. Our experiments revealed that mutations in *CAC1* or *RTT106* that directly deposit histones on the DNA, slowdown replication fork progression. In contrast, analysis of cells mutated in the intermediary *ASF1* or *RTT109* histone chaperones revealed that replisome progression is not affected. We found that mutations in histone chaperones including *ASF1* and *RTT109* lead to extended G2/M duration, elevated number of RPA foci and in some cases, increased spontaneous mutation rate. Our research suggests that histone chaperones have distinct roles in enabling high replisome progression and maintaining genome stability during cell cycle progression.

**Author Summary:** Histone chaperones (HC) play key roles in maintaining the chromatin structure during DNA replication in eukaryotic cells. Despite extensive studies on HCs, little is known regarding their importance for replication fork progression during S-phase. Here, we utilized a live-cell imaging approach to measure the progression rates of single replication forks in individual yeast cells mutated in key histone chaperones. Using this approach, we show that mutations in *CAC1* or *RTT106* HCs that directly deposit histones on the DNA lead to slowdown of replication fork progression. In contrast, mutations in *ASF1* or *RTT109* HCs that transfers H3/H4 to *CAC1* or *RTT106*, do not affect replisome progression but lead to post replication defects. Our results reveal distinct functions of HCs in replication fork progression and maintaining genome stability.

## Introduction

DNA replication in eukaryotic cells is a complex process that requires the accurate copying of the DNA itself and the formation of a precise chromatin structure [1,2]. The basic unit of chromatin is the nucleosome, composed of ~146 base pairs of DNA wrapped around an octamer of histones. A nucleosome is composed of a core of (H3-H4)_2_ tetramer and two flanking H2A-H2B dimmers [3,4]. During DNA replication, nucleosomes must be disassembled to allow replication fork progression and subsequently must be reassembled to establish the accurate chromatin state. Histone chaperones are essential for the process of DNA replication-coupled (RC) nucleosome deposition by facilitating correct histone assembly, post-translational modifications and localization during DNA replication [5–8].

Newly synthesized histones are assembled into nucleosomes by several histone chaperones including Asf1, Rtt109, Rtt106, Caf1 and the FACT complex that act in a sequential and coordinated manner to facilitate nucleosome deposition during DNA replication [5,9–11]. In budding yeast, Asf1 binds newly synthesized H3-H4 histones and promotes H3K56 acetylation by Rtt109 [12]. Acetylation of H3K56 enhances H3-H4 binding to Caf1 and Rtt106 that deposit (H3/H4)_2_ tetramers directly onto the DNA [13–15]. Previous studies have shown that single- or double-mutations of histone chaperone genes including Caf1 subunits, *RTT106, ASF1* or *RTT109* can lead to severe replication stress, checkpoint activation, increased recombination and sensitivity to DNA damaging agents [14,16,17]. However, the effect of histone chaperone mutations on the rate of replisome progression is much less understood. While studies in human cells have shown that mutations in genes essential for the synthesis of histones significantly slowdown replisome progression [18], studies in yeast show that mutations in histone chaperones can have little effect on S-phase progression [14,17,19–21].

One of the most important and extensively studied histone chaperone complex for RC nucleosome deposition both *in vitro* and in cells is the Caf1 complex [13]. Caf1 is composed of three subunits, Cac1, Cac2 and Cac3, which are highly conserved from yeast to human [22]. Cac1 is the largest subunit of the complex and was shown to interact with Proliferating Cell Nuclear Antigen (PCNA) through a canonical PCNA Interacting Protein (PIP) motif [23]. Since PCNA is an essential subunit of the replisome, Cac1-PCNA interaction can couple between nucleosome deposition and DNA synthesis [24]. In addition, Cac1 was shown to contain a Winged Helix Domain (WHD) motif allowing its direct binding to DNA [25]. While several biochemical studies provide important insights into the function of Caf1 subunits and the PIP and WHD domains of Cac1 [26,27], much less is known regarding their importance for replisome progression during S-phase and for nucleosome deposition on newly replicated DNA.

In this research, we have investigated the effect of deletion and/or mutations of major H3/H4 histone chaperones on DNA replication rate and G2/M duration. We have utilized our recently described live-cell microscopy approach [28,29] for the direct measurement of replisome progression and G2/M duration in individual live yeast cells (see below for details). We found that deletion of *CAC1* or *RTT106,* which directly deposit nucleosomes on the DNA, lead to slowdown of replication fork progression. Further analysis of point mutations in Cac1 showed a separation of function in Cac1 WHD and PIP domains affecting replisome progression and G2/M duration, respectively. In addition, we found that deletion of *ASF1* or *RTT109* histone chaperones that transfer H3/H4 histones to *CAC1* and *RTT106* did not lead to slowdown of replication fork progression but led to severe post-replication defects. Cell cycle analysis of the histone chaperone mutant cells, indicated a significant elongation of G2/M duration, ssDNA accumulation and, in some cases, elevated spontaneous mutation rates. These results demonstrate that histone chaperones exhibit distinct roles in facilitating high replisome progression rate and maintaining genome stability enabling efficient cell cycle progression.

## Results

### The experimental approach for measuring replication fork progression and G2/M duration in histone chaperone mutant strains

In order to examine the importance of Caf1, Rtt106, Asf1 or Rtt109 for replication fork progression during S-phase, we directly measured DNA replication rates in live WT and mutant cells [28,29]. Recently, we have described a live cell imaging approach for measuring the progression rates of single DNA replication forks in individual yeast cells [28]. This approach is based on the site-specific integration of arrays of *lacO* and *tetO* bacterial operator sequences, bound by the respective GFP-lacI and tetR-tdTomato cognate repressors, allowing the labelling of specific chromosomal loci as two distinct fluorescent dots (**Fig. 1**). The fluorescent intensity of these dots increases during replication of the *lacO* and *tetO* arrays since more fluorescently labelled repressor proteins are recruited to the newly replicated arrays (**Fig. 1**). These arrays are utilized as reporters of replication fork progression by inserting one array adjacent to an early origin of replication and the other array downstream in the same replicon (**Fig. 1**). By monitoring replicating cells and quantifying the intensity of each dot over time using live-cell fluorescence microscopy, the replication time of each array region is determined. Since the distance between the arrays along the chromosome is known, fork progression rate can be derived. We recently demonstrated the applicability of this approach by examining replication at different loci, in the presence of different concentrations of HU and in different mutant strains [28]. More recently, we have utilized this approach to examine replication fork progression through G-quadruplex structures on the background of different *pif1*-mutant strains [29]. An additional advantage of this system is the ability to monitor yeast cell morphology and detect anaphase events at the single-cell level. Anaphase can be monitored as splitting of each fluorescent dot into two, one of which moves to the daughter cell. By calculating the time interval between replication of the second array (*tetO*) and anaphase, we can estimate the duration of G2/M and examine how it is affected in different histone chaperone mutant cells (**Fig. 1**) [28].

**Figure 1:**
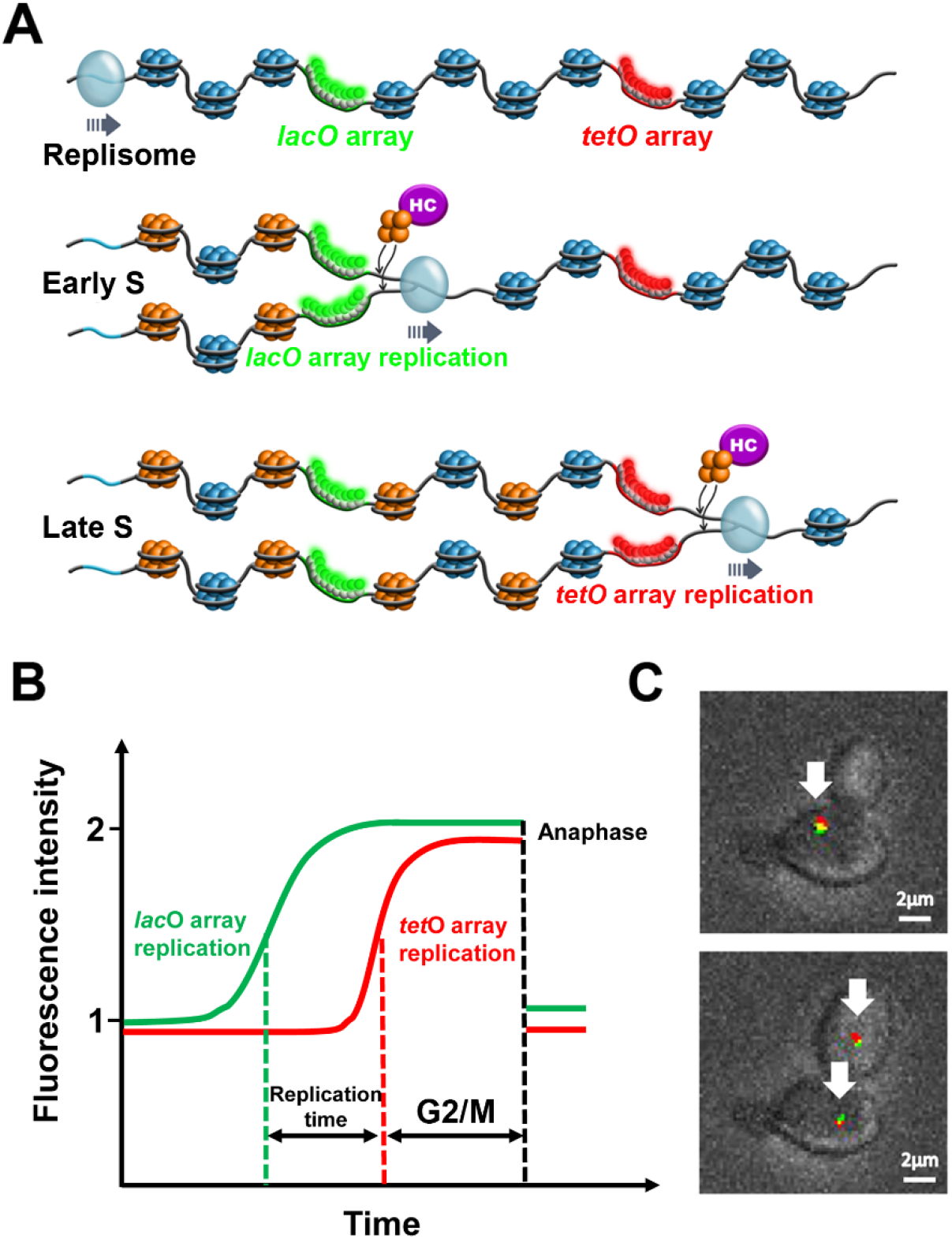
Measuring replication rates in histone chaperone mutant yeast cells. (**A**) Schematic illustration of the design for the real-time analysis of replication kinetics in live yeast cells containing mutations in histone chaperone genes. Strains are genetically engineered to contain *lacO* and *tetO* arrays located 20 Kb apart, downstream from an origin of replication. Binding of GFP-lacI and tetR-tdTomato lead to green and red fluorescent foci, respectively. During DNA replication, array duplication leads to recruitment of additional lacI-GFP and tetR-tdTomato proteins leading to an increase in fluorescence intensity. The newly-synthesized DNA is packed around new (orange) and pre-existing (blue) nucleosomes. Histone chaperones (HC, purple) deposit newly-synthesized histones as (H3/H4)_2_ tetramers behind the replisome. The replication time of each locus is measured using time-lapse confocal microscopy. For simplicity, origin firing is shown only to the array direction. (**B**) Schematic display of the doubling of fluorescent intensity of the GFP-lacI and tetR-tdTomato foci in a single cell due to *lacO* and *tetO* array duplication during DNA replication. The time between replication of the arrays (replication time) is calculated using the mid-rise points of the GFP and tdTomato fluorescence intensities. The G2/M duration is estimated by the time delay between the mid-rise point of the tdTomato fluorescence intensity and the anaphase (**C**) Image of single cells from the yeast strains used in this study. Top - image following replication before anaphase, Bottom – after anaphase. Images scale: 2μm

### The importance of Caf1 complex subunits for replisome progression and G2/M duration

Previous studies have shown that Caf1 complex, composed of Cac1, Cac2 and Cac3, is important for H3/H4 deposition directly onto the DNA[13,27]. To examine the importance of each subunit of Caf1 complex for replisome progression rate and G2/M duration, we have deleted *CAC1, CAC2* or *CAC3* subunits on the background of a strain containing the *lacO* and *tetO* arrays located at the vicinity of ARS413. We then measured replication times and G2/M duration in individual yeast cells using live cell microscopy as described above (**Fig. 2AB** and **Table S1**). We found that *cac1*- deletion leads to a significant elongation of replication times relative to the WT strain while *cac2*- or cac3-deletion do not (**Fig. 2C**). Next, we analyzed the G2/M duration in the different strains and found that *cac1*-deletion cells exhibit significant increase in G2/M duration relative to WT cells (**Fig. 2D**). These results highlight the importance of Cac1 for replisome and G2/M progression and are in good agreement with a previous study showing that Cac1 is essential for the Caf1 protein complex assembly and H3/H4 binding [13,26].

**Figure 2:**
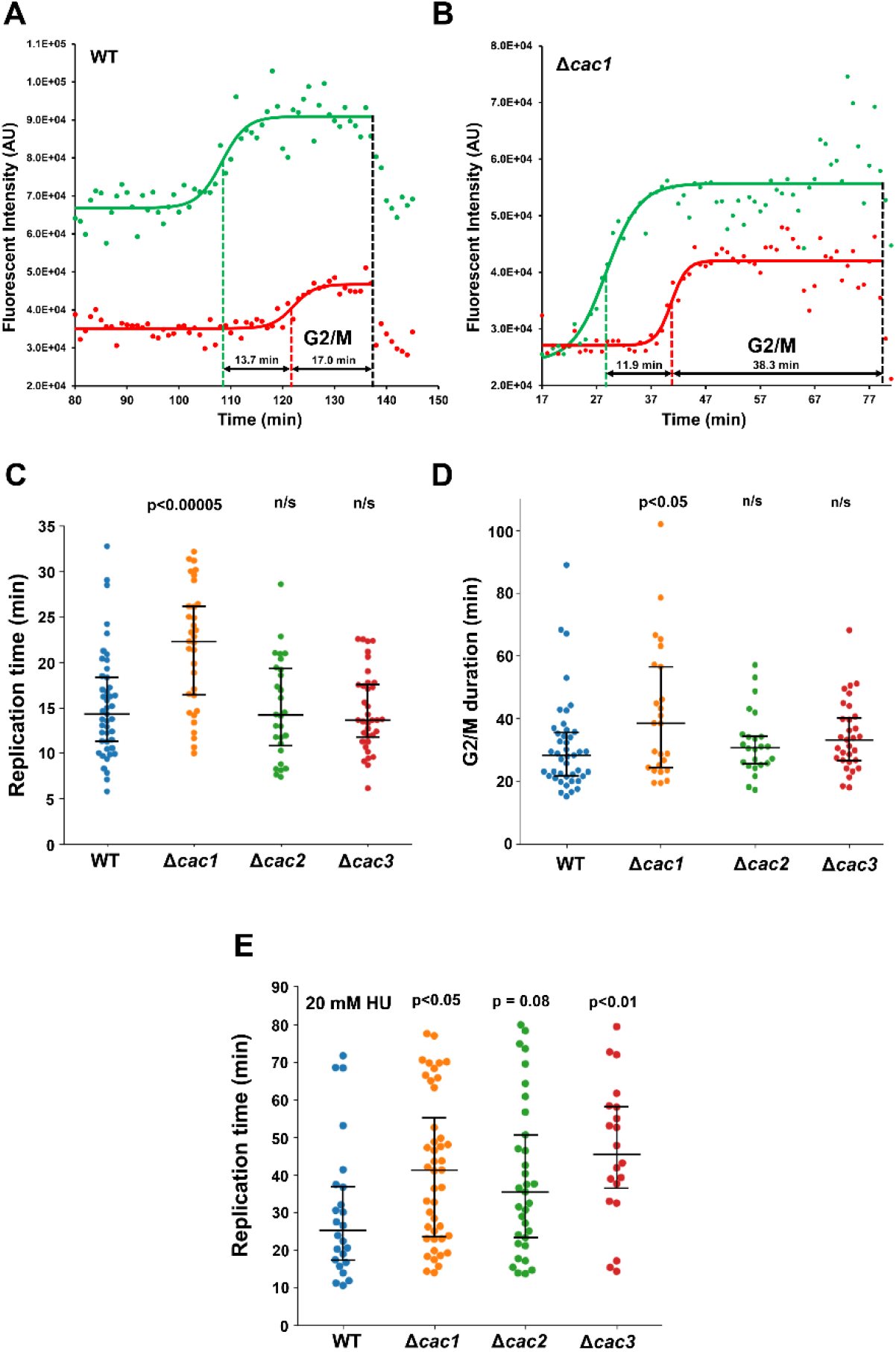
Extended replication times and G2/M duration in *cac1*-deleted cells. (**A-B**) Representative results of a single cell from (**A**) WT and (**B**) *cac1*-deleted strains showing a significant increase in G2/M duration in the *cac1-deleted* cell. Solid lines represent a fit of the data to a sigmoidal function showing the *lacO* array replication (green) and the *tetO* array replication (red), green and red mid-points are indicated by dashed lines while black dashed lines show the anaphase timepoint. Black double arrows measuring the time delay between *lacO* array replication and *tetO* array replication and *tetO* array replication and anaphase indicates the replication time and G/M duration, respectively. (**C**) Replication times of ~30.6 Kb at the vicinity of ARS413 for WT, *cac1*-, *cac2*- and *cac3*-deleted cells. (**D**) G2/M durations, estimated by measuring the time delay between the mid-rise point of the tdTomato fluorescence intensity and the anaphase, for WT, *cac1-, cac2-* and cac3-deleted cells. (*E*) Replications times as in (**C**) but in the presence of 20 mM of HU. Significance was determined by Monte Carlo resampling and p values relative to WT are shown. The number of cells analyzed for each strain, all median values and statistical results are shown in **Table S1** and for (**E**) **Table S2.**

Previous studies have revealed the contribution of the Caf1 subunits for H3/H4 assembly and nucleosome deposition under DNA stress conditions [30–32]. To further investigate the effect of *cac2-* or cac3-deletion on DNA replication under these conditions, we incubated the Caf1 mutant strains with 20 mM HU and measured their replication times as described above. We found that all Caf1 component mutants exhibited longer replication times relative to WT (**Fig. 2E** and **Table S2**). These results are in agreement with previous studies [30–32] and deepen our understanding about the importance of Caf1 subunits on DNA replication under stress conditions.

### G2/M elongation due to *cac1*-deletion is linked to spindle checkpoint activation

G2/M elongation may stem from the activation of two mitotic checkpoints due to genomic instability accumulated during DNA replication [33]. These two checkpoints are the DNA damage checkpoint and the spindle assembly checkpoint [34,35]. Previously, mutation in *CAC1* was shown to be implicated in the activation of the spindle checkpoint [36]. Specifically, it was shown that the deletion of *MAD2,* the master checkpoint protein of spindle assembly, suppressed the mitotic delay measured in *cac1*- and *hir1-double* deletion strain [36]. To examine whether the G2/M elongation observed in cac1-deletion cells (**Fig. 2**) was due to spindle checkpoint arrest, we deleted *MAD2* in the background of *cac1*- deletion. We then measured replication times and G2/M duration in these mutant cells using the live-cell imaging system described above. In accordance with the previous study [36], we found that the elongation of G2/M duration due to *CAC1* deletion is suppressed in the double mutant cells and is similar to the G2/M duration values measured for WT cells (**Fig. 3**). To further support the importance of spindle checkpoint activation in cac1-deleted cells, we examined the growth of *cac1-* and *mad2-double* deleted cells using a spot assay. In agreement with the microscopy experiments, we found that the addition of *MAD2* deletion suppressed the growth defects observed in cac1-deleted cells (**Fig. S1**). These results, further supports the role of *CAC1* in ensuring mitotic spindle integrity prior to cell mitosis.

**Figure 3:**
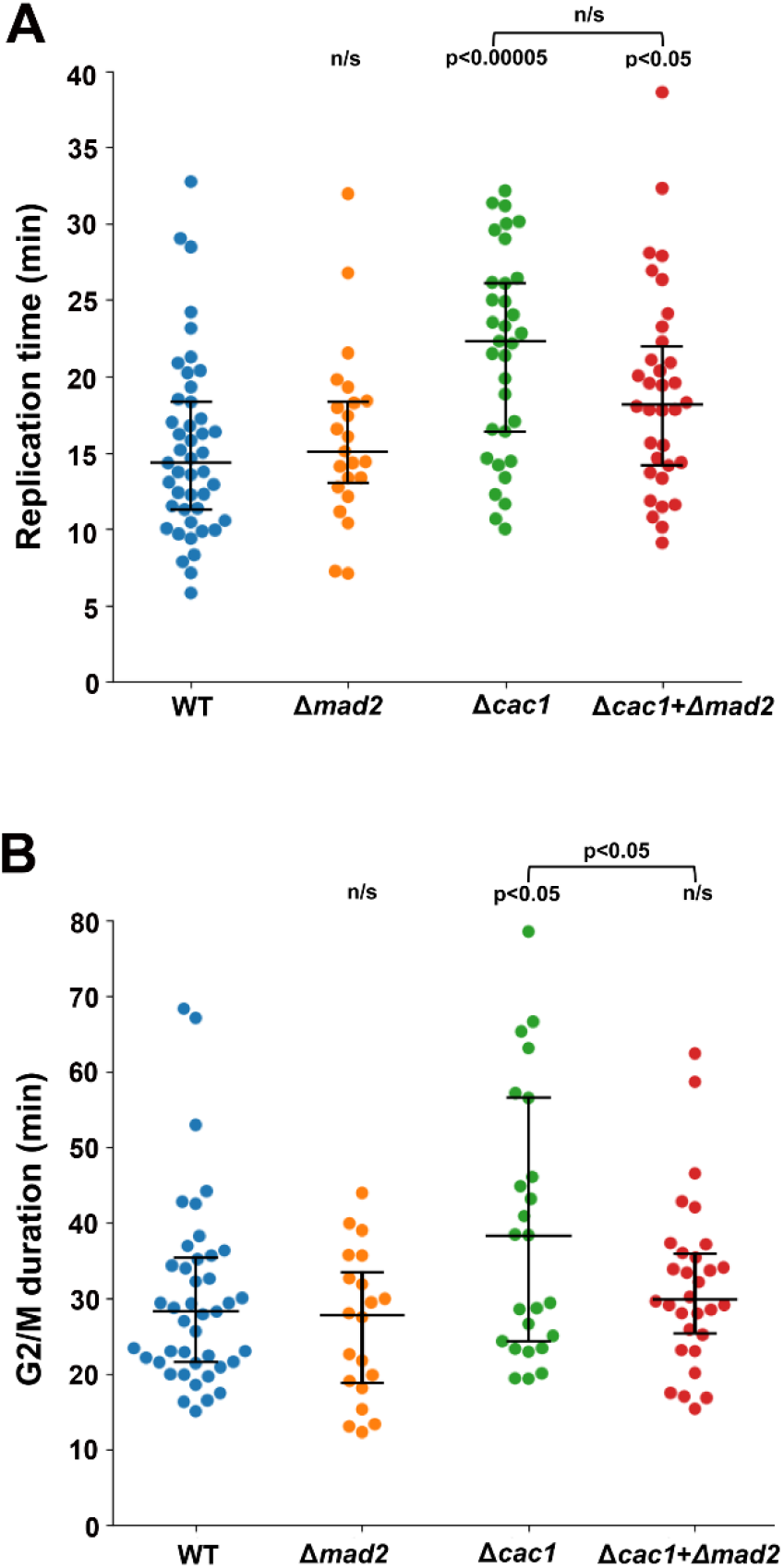
Extended G2/M duration in *cac1*-deleted cells is suppressed by the deletion of *MAD2:* (**A**) Replication times of ~30.6 Kb at the vicinity of ARS413 for WT, *mad2*-, *cac1*- deletion and *mad2* and *cac1*-double deleted cells. (**B**) G2/M durations, estimated by measuring the time delay between the mid-rise point of the tdTomato fluorescence intensity and the anaphase, for WT, *mad2-,* cac1-deletion and *mad2* and cac1-double deleted cells. The number of cells analyzed for each strain and all median values and statistical results are shown in **Table S1.**

### Separation of function of the WHD and PIP regions in Cac1

The elongation of replication times and G2/M duration of cac1-deleted cells can stem from the lack of Caf1 complex formation or from deficiencies in Cac1 specific interactions. Previously, Cac1 was shown to contain a WHD domain and a PIP region facilitating its direct interaction with the DNA and PCNA, respectively [23,25]. To examine the importance of these Cac1 regions for replisome progression and G2/M duration, we have generated strains for replication measurements containing K564E/K568E and F233A/F234G mutations in Cac1 WHD domain and the PIP region, respectively. These mutations in the WHD and PIP domains of Cac1 were previously shown to abolish binding to DNA and PCNA, respectively [23,25]. Next, we measured replication time and G2/M duration in individual cells containing these mutations as described above. We found that mutating Cac1 WHD domain leads to increased replication times with no effect on the G2/M duration, relative to WT cells (**Fig. 4A**). In contrast, we found that mutating the PIP region of the protein does not affect replication times but leads to a significant elongation of G2/M duration (**Fig. 4B**). These results demonstrate the separation of function between the WHD and PIP regions of Cac1, with DNA binding contributing to proper replisome progression and PCNA binding preventing replication stress.

**Figure 4:**
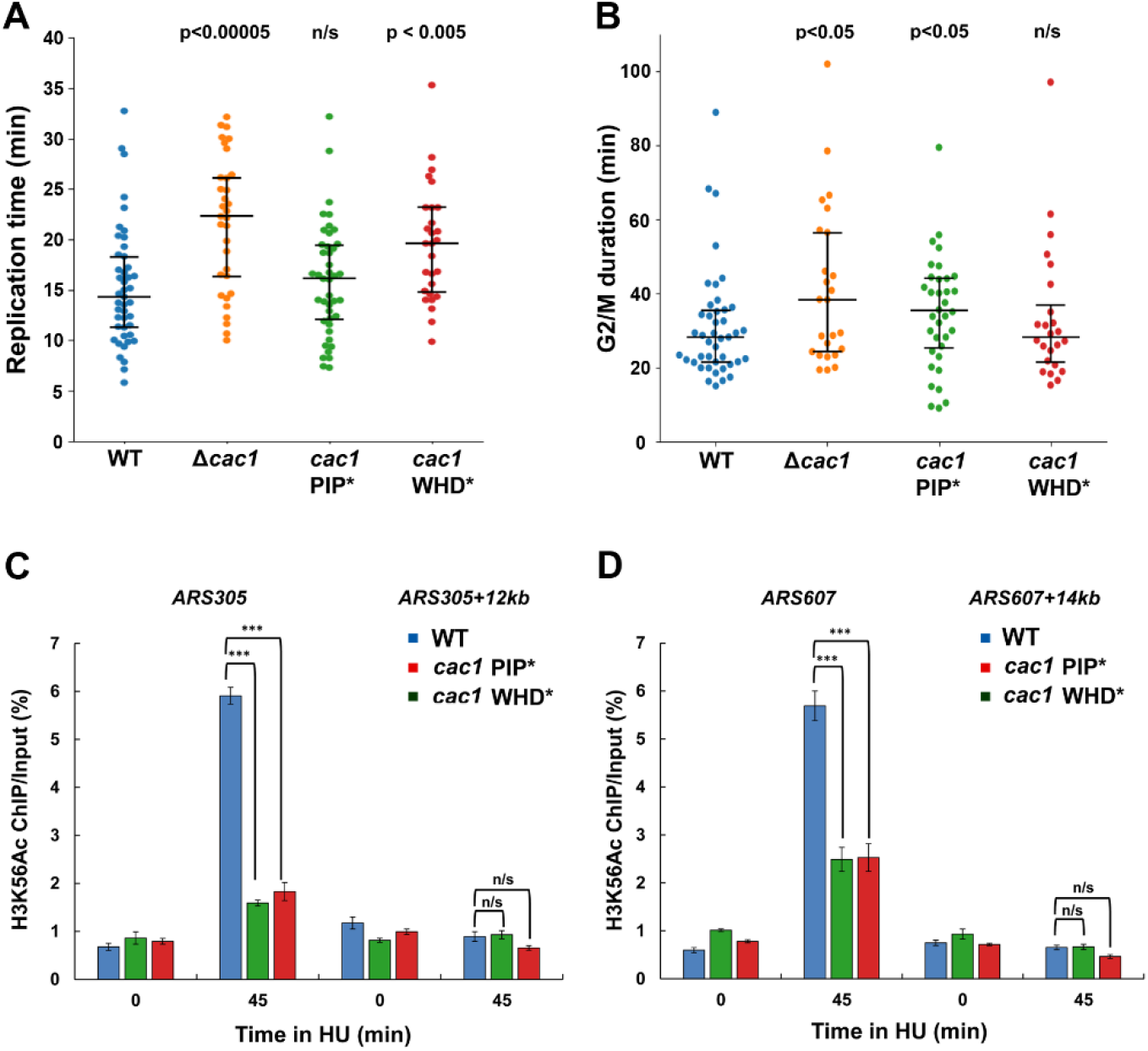
Replication times, G2/M duration and ChIP analysis of *cac1* mutated cells. Mutations in the Cac1 WHD an PIP domains lead to increased replication times and G2/M duration, respectively, and lower nucleosome deposition during DNA replication, relative to WT cells. (**A**) Replication times of ~30.6 Kb at the vicinity of ARS413 for WT, *cac1*- deletion, *cac1* PIP* (PIP mutant containing the F233A and F234G mutations)- and *cac1* WHD* (WHD mutant containing the K564E and K568E mutations)-mutant cells. (B) G2/M durations, calculated as described in **Fig. 2D**, for WT, cac1-deleltion, *cac1* PIP*- and *cac1* WHD-mutant cells. Significance was determined by Monte Carlo resampling and p values relative to WT are shown. The number of cells analyzed for each strain and all median values and statistical results are shown in **Table S1. (C-D).** ChIP analysis of H3K56Ac deposition for WT and *cac1* mutant strains. WT and *cac1* mutant cells were arrested at G1 phase using α-factor at 25°C and then released into fresh medium containing 0.2 M HU. Equal numbers of cells were collected just prior to (G1, 0 min) and at 45 minutes following release into fresh medium containing HU. ChIP assays were performed using antibodies against H3K56Ac and H3 (as a normalization control). The ChIP DNAs were analyzed using four different primer pairs amplifying (**C**) *ARS305, ARS305+12kb* and (**D**) *ARS607, and ARS607+14kb,* using quantitative PCR. At least three independent biological repeats were performed, with similar trends in all cases. The mean and SDs of three independent biological replicates are shown. *P* values derived from two-way analysis of variance (ANOVA) (*\*\*\*P* value ≤ 0.001). ChIP over input is shown here, results were similar after normalizing to total H3 ChIP for the same trend (**Fig. S2**), indicating that loss of H3K56Ac signal is not as a result of decreased total H3.

Next, we examined the level of nucleosome deposition on newly synthesized DNA in the WHD and PIP Cac1 mutant strains at the vicinity of ARS305 and ARS605 using chromatin immunoprecipitation (ChIP) following release from HU arrest. In accordance with the effect of these mutations on replisome progression rate and G2/M duration, we found a significant decrease in the level of nucleosome deposition during DNA replication in these mutant strains relative to the WT (**Fig. 4C-D** and **Fig. S2**). To further examine the effect of HU replication stress on the two Cac1 mutants, we measured their replication times in presence of 20 mM HU. We found that both mutants exhibit more than 9 minutes longer replication times compared to WT (**Fig. S3**). These results highlight the importance of both PIP and WHD of Cac1 for nucleosome deposition and replication fork progression at early S-phase under DNA stress conditions.

### Analysis of mutation in *RTT106*, *RTT109* or *ASF1* histone chaperones

To examine how mutations in additional H3/H4 histone chaperones affect replisome progression and G2/M duration, we have generated deletions in *RTT106, RTT109* or *ASF1* on the background of the *lacO* and *tetO* containing strain as described above. In addition, to examine how double deletion in *CAC1* and *RTT106* affect these properties we have generated this strain on the same background. Rtt106 was previously shown to interact with H3/H4 and the DNA to enable (H3/H4)_2_ tetramer deposition on newly synthesized DNA [37]. Cac1 and Rtt106 were shown to physically interact suggesting that these histone chaperones cooperate to facilitate high level of nucleosome deposition on newly synthesized DNA [38]. Asf1 and Rtt109 are additional key histone chaperone and histone acetylase, respectively, acting prior to Cac1 and Rtt106 to facilitate H3K56 acetylation of newly synthesized histones [12,19]. Analysis of *rtt106*-deletion and the *cac1-rtt106-double* deletion cells using our assay showed significantly increased replication times, relative to WT cells. In contrast, *rtt109*- or *asf1*-deletion did not lead to a significant change in replication times (**Fig. 5A**). Analysis of the G2/M duration in *cac1*- rtt106-double deletion, *rtt109-* or *asf1*-deleted cells was significantly extended while was not affected in *rtt106*-deleted cells relative to WT cells (**Fig. 5B**). Elongation of G2/M duration, observed in *cac1-* rtt106-double deletion, *rtt109*- or *asf1*-deleted cells, is in agreement with several studies showing checkpoint activation and replication stress in these strains [14,19–21].

**Figure 5:**
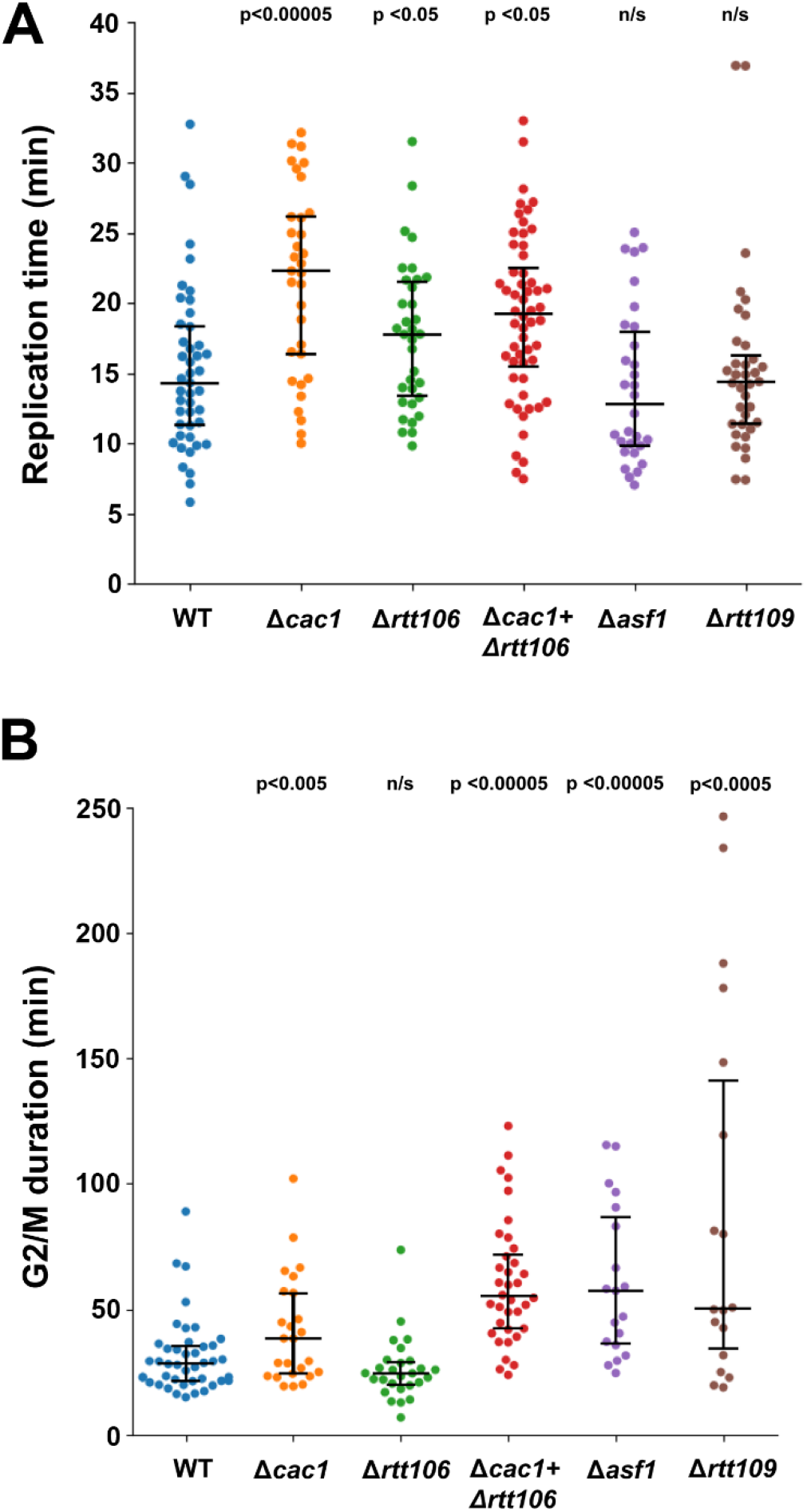
Replication times and G2/M durations of *cac1*-, *rtt106*-, *cac1*- and *rtt106*-, *asf1*- or *rtt109*-deleted cells. Cell measurements are shown as swarm plots (**A**) Replication times of ~30.6 Kb at the vicinity of ARS413 for WT, *cac1*-, *rtt106*-, *cac1*- and *rtt106*-, *asf1*- or *rtt109*-deleted cells. WT and *cac1*-deleted cells are shown as comparison. (**B**) G2/M durations, estimated by measuring the time delay between the mid-rise point of the tdTomato fluorescence intensity and the anaphase, for WT, *cac1* -, *rtt106-,cac1-* and *rtt106-, asf1-* or *rtt109*-deleted cells. Significance was determined by Monte Carlo resampling and p values relative to WT are shown. The number of cells analyzed for each strain and all median values and statistical results are shown in **Table S1.**

### Increased G2/M duration in histone chaperone mutant cells is associated with ssDNA accumulation

Replication stress often lead to the accumulation of single-stranded DNA near active replication and repair sites [39]. This accumulation can be visualized as foci of the ssDNA-binding protein Replication Protein A (RPA) [40–42]. To examine whether long G2/M duration in histone chaperone mutant cells is associated with increased level of ssDNA accumulation, we monitored the levels of RPA foci in these mutant cells. For monitoring RPA foci in live cells, we used Rfa1, expressed under its native promoter, tagged with GFP at its N-terminus [43] and used fluorescent time-lapse microscopy to monitor foci in cells that started budding. To verify the nuclear localization of the RPA foci, we monitored their co-localization with a nuclear signal generated by simultaneous expression of NLS-tetR-tdTomato. Indeed, we found that RPA foci are identified within the cell nucleus following budding and entering S-phase (**Fig. S4**). Previously, the lack of PCNA ubiquitylation by Rad18 ubiquitin ligase was shown to significantly increase the level of RPA foci in mammalian cells [42]. Thus, to further validate our system, we generated *rad18*- deletion strain and examined the level of RPA foci in mutant cells relative to WT. We found that the percentage of cells with RPA foci containing the rad18-deletion is significantly higher than WT cells (**Fig. S5**).

Next, we generated the different histone chaperone mutants on the background of GFP-Rfa1 strain and monitored the accumulation of RPA foci in mutant and WT cells (please see **Fig. 6A** for representative WT and *asf1*-deleted cells). We found an increase in the percentage of cells containing RPA foci in the histone chaperone mutant cells relative to WT cells indicating that these mutations lead to the accumulation of ssDNA [39,40] (**Fig. 6B**). Increase in RPA foci was most prominent in *asf1*- or *rtt109*-deleted strains, in which over 40% of the cell population contained at least one clear RPA focus. We found that the increased level of RPA foci in histone chaperone mutant strains is strongly correlated with increased G2/M duration, validating that the delayed G2/M phenotype seen in these strains is related to accumulation of DNA damage during S phase (**Fig. S6**). Taken together, our results suggest that replication that is uncoupled to nucleosome deposition can lead to replication stress, accumulation of ssDNA, and significant delays in cell cycle progression (**Fig. 6** and **Fig. S6**).

**Figure 6:**
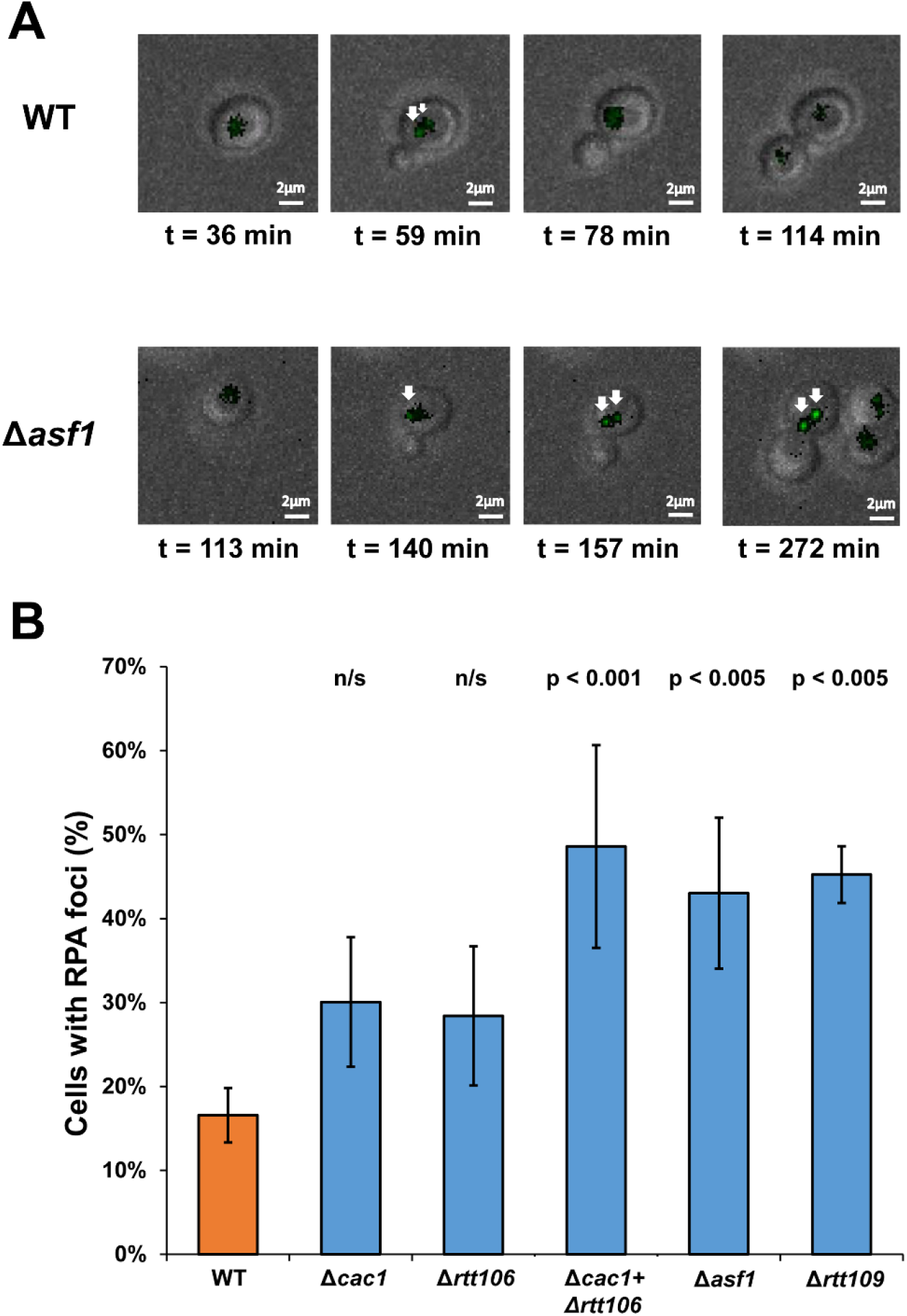
RPA foci accumulate in histone chaperone mutant cells. (**A**) Representative WT and *asf1*-deleted cells containing RPA foci are shown at different time points. In the case of the WT cells, the RPA focus disappears in late S-phase. In the case of *asf1*-deleted cell, mitosis does not take place even after 120 min from initial budding and the RPA foci persists. (**B**) WT or histone chaperone mutant cells expressing N-terminal GFP-Rfa1 fusion were analyzed using time-lapse fluorescent microscopy. At least 60 cells from three independent experiments were analyzed for each strain and scored for the presence of one or more RPA foci during S-phase. Images scale: 2μm

### Spontaneous mutation rate analysis of histone chaperone mutant strains

Replication fork stalling and accumulation of ssDNA during replication can lead to double strand breaks that can be detrimental to cell survival [44,45]. To avoid double strand breaks, cells employ a variety of mechanisms including the recruitment of translesion DNA polymerases (TLS) by the *RAD6/RAD18* pathway [46,47]. This pathway, initiated by mono-ubiquitylation of PCNA on K164, can minimize fork stalling but may lead to increased mutation rates due to the inaccuracy of the TLS polymerases [46,47]. To examine whether mutation rates are elevated in the different histone chaperone mutant strains, we have utilized the *CAN1* assay. We have previously utilized this assay to examine a variety of PCNA mutants leading to *RAD6/RAD18* pathway activation [48]. In accordance with previous analysis, we found that *asf1-* or rtt109-single deletions lead to a mild increase in mutation rate of up to two-fold relative to the WT strain [49]. Single deletions of *CAC1* or *RTT106* lead to similarly mild increase. In contrast, we found that the *cac1-rtt106-double* deletion leads to a dramatic increase of ~7 fold in mutation rate, (**Fig. 7** and Table 1). To examine whether the increased mutation rate in the *cac1-rtt106-double* deletion strain is due to *RAD6/RAD18* pathway activation, we generated a *cac1-rtt106-rad18-triple* deletion strain and examined its mutation rate. Consistent with *RAD6/RAD18* pathway activation, we observed that the triple deletion strain exhibits a dramatic reduction in mutation rate relative to the *cac1*-*rtt106*-double deletion strain (**Fig. 7**). In addition, we found that depletion of DNA polymerase ζ, a key TLS polymerase, in this strain leads to higher sensitivity to MMS, highlighting its possible activation upon DNA damage (**Fig. S7**). Taken together, our results with the *cac1-rtt106-double* deletion strain show that mutations in histone chaperones can lead to *RAD6/RAD18* pathway activation and increased spontaneous mutation rates. However, results with the other histone chaperone mutant strains, exhibiting long G2/M duration and a mild increase in mutation rate (e.g. *rtt109* or *asf1* mutant strains), suggest that other pathways can be activated upon ssDNA accumulation during DNA replication (see Discussion section).

**Figure 7:**
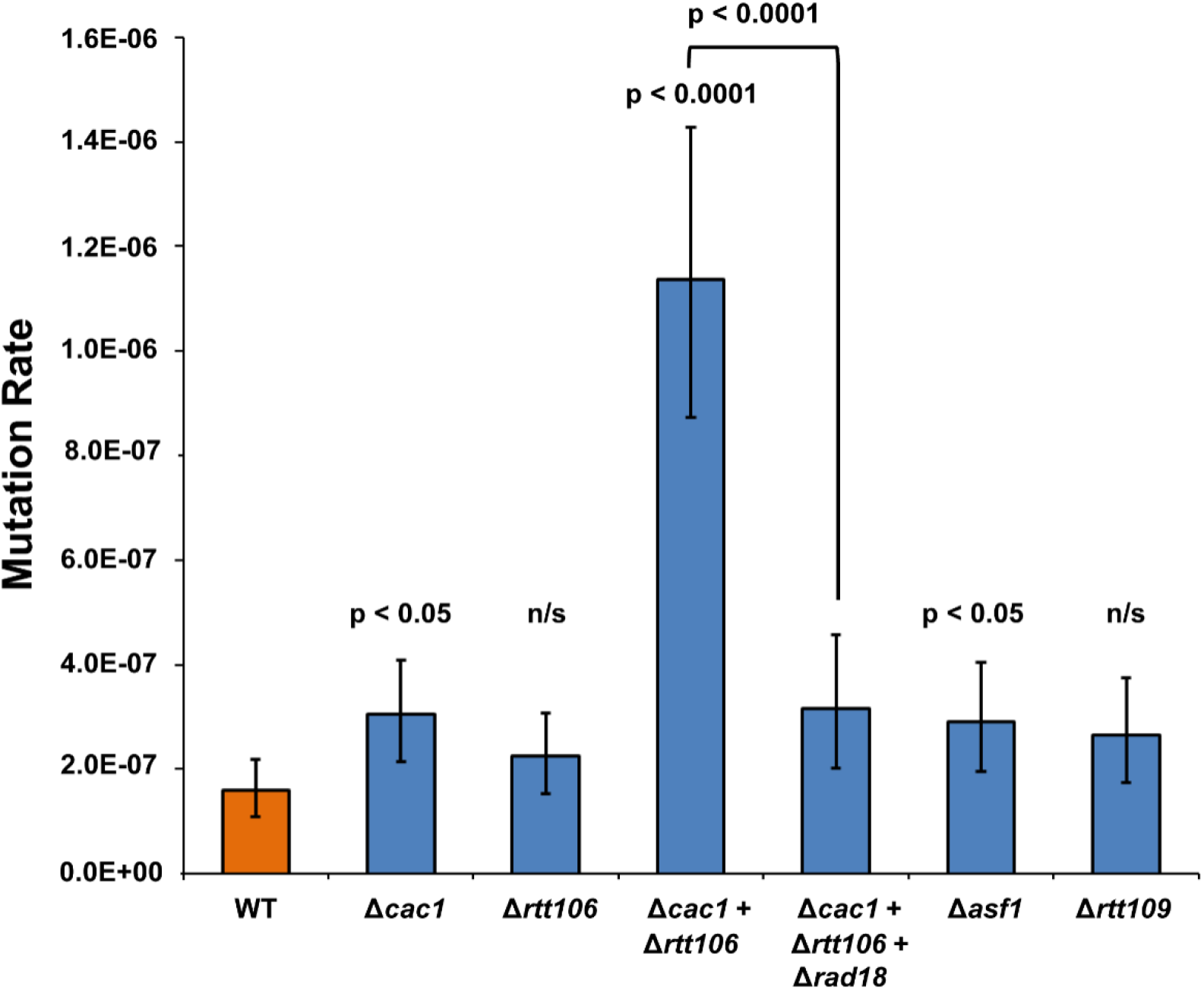
Increased spontaneous mutation rate in histone chaperone mutants is most significant in the *cac1*-*rtt106*-double deletion strain. Spontaneous mutations in this strain are dependent upon *RAD6/RAD18* pathway since spontaneous mutation rate in the triple *cac1*-*rtt106*-*rad18*-deletion strain is dramatically reduced relative to the *cac1-rtt106-double* deletion strain. The data are rates of Can^r^ mutation values within the 95% confidence limits of at least 16 independent analyses. The values of the spontaneous mutation rate of the different strains are presented in **Table 1.**

**Table 1:** Spontaneous mutation rate at the *CAN1* locus

| Genotype | Mutation rate Can<sup>R</sup> [CI]<sup>a</sup> | Fold increase compared to WT |
| --- | --- | --- |
| WT | 1.59 [1.09, 2.19] x 10<sup>-7</sup> | - |
| $\Delta cac1$ | 3.04 [2.14, 4.08] x 10<sup>-7</sup> | 1.9 |
| $\Delta rtt106$ | 2.24 [1.54, 3.08] x 10<sup>-7</sup> | 1.4<sup>b</sup> |
| $\Delta cac1 + \Delta rtt106$ | 11.4 [8.72, 14.30] x 10<sup>-7</sup> | 7.2 |
| $\Delta cac1 + \Delta rtt106 + \Delta rad18$ | 3.15 [2.01, 4.57] x 10<sup>-7</sup> | 2.0 |
| $\Delta asf1$ | 2.90 [1.95, 4.05] x 10<sup>-7</sup> | 1.8 |
| $\Delta rtt109$ | 2.64 [1.75, 3.74] x 10<sup>-7</sup> | 1.7<sup>b</sup> |

<sup>a</sup>The numbers in brackets represent the low and high values for the 95% confidence interval for each rate, obtained using the confidence interval for the median test. The medians and 95% confidence intervals were deduced from at least 16 independent determinations for each strain.
<sup>b</sup>Non significant difference from the WT

## Discussion

DNA RC nucleosome assembly that connects DNA synthesis with nucleosome deposition and modification has been extensively studied in the past two decades [9,11,50]. These studies allowed the identification and characterization of histone chaperones that are important for nucleosome deposition on newly synthesized DNA. However, much less is known regarding how mutations in histone chaperones affect replisome progression during DNA replication. Here, we used our recently described direct approach to monitor replisome progression in individual yeast cells [28,29] that are mutated in different histone chaperone genes, including Caf1 subunits, *RTT106, ASF1* and *RTT109.* We found that mutation in Cac1 or Rtt106 that directly deposit H3/H4 histones on DNA, significantly slowdown replication fork progression (**Fig. 2**–**5**). These results suggest that the direct interaction of these chaperones with the replication fork is essential for enabling high fork progression rate during S-phase. Interestingly, we found that mutations in histone chaperones which transfer H3/H4 on Cac1 or Rtt106 prior to their final assembly, including *ASF1* or *RTT109* histone acetyl transferase, do not affect replisome progression but significantly extend G2/M duration (**Fig. 5**) highlighting the distinct and independent roles of histone chaperones in genome replication and stability. The normal replisome progression in *asf1*-deleted cells (**Fig. 5**) is in good agreement with previous studies showing that yeast budding and progression through S-phase is not significantly altered in this mutant strain [20,21].

Our system also allows the detection of elongations in G2/M duration by monitoring both fluorescence and morphological changes in individual cells. Our findings that single deletion of *CAC1, RTT109* or *ASF1* or double deletion of *CAC1* and *RTT106* (**Fig. 5**) leads to a significant increase in G2/M duration highlight the important role of these chaperones in preventing replication stress [39]. Our analysis of G2/M duration is probably an underestimation of the true G2/M length in these mutant cells since high percentage of the analyzed cells did not undergo mitosis during the experiment duration (Table S3). Our further examination of these strains revealed a dramatic increase in RPA foci formation following DNA replication (**Fig. 6**). We found that in the case of double deletion of *CAC1* and *RTT106* the elongated G2/M duration and RPA foci formation is also correlated with a significant increase in spontaneous mutation rate (**Fig. 7**). Previously, several studies have extensively examined different aspects of genome stability in histone chaperone mutant strains. An extensive study examining several single and double deletions of histone chaperones including *RTT109, ASF1, CAC1* and *RTT106* in yeast revealed that most mutations lead to increased recombination frequency, checkpoint activation, loss of replisome stability and sensitivity to DNA damaging agents [14]. In addition, *RTT106* was shown to genetically interact with *DDC1-MEC3- RAD17* that forms an alternative sliding clamp termed the 9-1-1 complex which is involved in checkpoint activation [51]. Furthermore, *ASF1* was shown to directly interact with the downstream *RAD53* effector kinase [52,53] suggesting that these checkpoint proteins are intimately connected to DNA RC nucleosome deposition. Our findings that histone chaperone mutants lead to high level of RPA foci formation highlight their importance for preventing the accumulation of ssDNA. It complements studies showing the interaction between histone chaperones and S-phase checkpoint activation where ssDNA accumulation sensed by RPA (**Fig. 6**) is transmitted to several proteins including the 9-1-1 complex [51], leading to Mec1 sensor kinase activation followed by Rad53 effector kinase activation [52,53]. The correlation between high level of RPA foci formation and G2/M duration (**Fig. S6**) highlights the detrimental effect of ssDNA accumulation for cell cycle progression. In contrast to WT cells in which RPA foci can be corrected during S-phase, we observed that for histone chaperone mutants, RPA foci can be maintained through the G2 phase following DNA replication (**Fig. 6A**).

Analysis of the different subunits of the Caf1 complex indicated that deletion of *CAC1* exhibits the strongest effect on both replisome progression and G2/M duration (**Fig. 2**). Previously, Cac1 has been implicated in the MAD2-dependent mitotic spindle checkpoint [36] and our results further verify this study (**Fig. 3**). Further analysis of mutations in Cac1 WHD or PIP regions showed that while the WHD domain is important for replisome progression, the PIP region is important for preventing replication stress (**Fig. 4**). This separation of function highlights the importance of direct Cac1-DNA interaction, mediated by the WHD domain, for fast replisome progression. In contrast, elongation of G2/M duration in Cac1 PIP mutant highlights the importance of coordinating Cac1 activity with replisome progression for avoiding replication stress. Our further analysis of the WHD and PIP Cac1 mutants showed a significant reduction of nucleosome deposition of both mutants relative to WT strain as well as slowdown of replisome progression in the presence of HU (**Fig. 4** and **Fig. S3**). Previously, several studies have examined the importance of Cac1 WHD and PIP regions for their biochemical and cellular functions including their importance for DNA binding following association with H3-H4 histones [27], nucleosome assembly *in vitro* [23] and sensitivity to DNA damaging agents [25]. Our results examining the function of these regions in live cells are in agreement with these studies and further enhance our understanding of the contribution of each interaction for efficient replisome progression and short G2/M duration.

Previously, mutations in histone chaperones were shown to increase recombination level and gross chromosomal rearrangement highlighting their roles in maintaining genome stability [14,16]. Asf1, Caf1 and Rtt109 were shown to be important for deactivation of damage checkpoint after double strand break repair [55,56]. However, much less is known regarding their contribution to DNA damage response pathways that include the activation of TLS polymerases for damage bypass. Our findings that mutations in histone chaperones lead to moderate increase in spontaneous mutation rate (**Fig. 7**) suggest that other pathways are dominant, including homologues recombination, to overcome replication stress. Our findings are in good correlation with a previous study examining the level of spontaneous mutation rate in several deletion strains including the *RTT109* or *ASF1* single deletions [49]. The significant increase in spontaneous mutation rate in cells deleted in *CAC1* and *RTT106* (**Fig. 7**) may suggest that these chaperones link PCNA both to nucleosome deposition and activation of TLS polymerases through PCNA ubiquitylation by the *RAD6/RAD18* pathway. Notably, these results are in agreement with the involvement of *CAC1* in the *RAD6/18* post-replication repair pathway [32].

Overall, our experiments reveal that histone chaperones exhibit distinct functions in replisome progression and maintaining genome stability. Our data showing the effects of *cac1-or* rtt106-deletion on replisome progression highlight the possible importance of direct interaction of these chaperones with H3/H4 and the DNA in facilitating fast replisome progression. In contrast, histone chaperones which are involved indirectly with the transfer of histones on DNA, including Asf1 and Rtt109 do not significantly affect replisome progression but are essential for preventing replication stress. Our system can be further utilized to investigate how additional histone chaperones, such as the FACT complex, Nap1, Vps75 [5,57], or combinations of mutants affect replication fork progression and G2/M duration.

## Materials and Methods

### Strain generation

Strains for replication time measurement were generated on the background of a W303 MATa *Saccharomyces cerevisiae* strain, expressing GFP-LacI and tetR-tdTomato fusion proteins in the nucleus. *lacOx256* and tetOx224 arrays are located at chrIV:332960 and chrIV:352560 respectively, near ARS413 and with a mid-array distance of approximately 30.6 kb [28]. *CAC1, CAC2,* CAC3, *RTT106, ASF1, RTT109* and *MAD2* genes were replaced by *natMX* or *hphMX* antibiotic cassettes using the Lithium Acetate (LiAc) transformation method [58]. The *cac1* WHD, *cac1* PIP, and *rtt106* KK245,246AA mutants were generated on the background of the *cac1-* or *rtt106-* deleted cells respectively, with a markerless CRISPR/Cas9 approach, by targeting either the *natMX* or *hphMX* with specific gRNAs and the respective mutant genes as DNA donors [59]. All replacements were validated with Polymerase Chain Reaction (PCR) and Sanger sequencing.

### Microscopy and data analysis

All microscope experiments for the determination of replication time were conducted as previously described [28] with slight modifications. Briefly, yeast cultures were grown overnight to OD_600nm_ = 0.1-0.2 in synthetic complete (SC) medium containing 4% glucose at 30°C. The cultures were then synchronized at G1 using 10 μg/ml α-factor (GenScript) for 2-3 hours. For hydroxyurea (HU) experiments, yeast cultures were incubated with 20 mM HU one hour before and during the imaging. Cells were then immobilized on microscopy slide chambers (Ibidi) coated with 2 mg/ml concanavalin A (Sigma-Aldrich) and washed thoroughly from α-factor with warm SC medium containing 4% glucose prior to microscopy measurements. Live cell imaging of the cells was performed on a Marianas spinning-disk confocal microscope (3i) using an Evolve EM-CCD camera (Photometrics) with 1 min intervals for 2-8 hours at 28°C, using a x63 oil objective (NA = 1.4) in 3D (12 z-sections at 0.7μm apart). GFP-LacI and TetR-tdTomato were excited with 488nm and 561nm lasers, respectively. For data analysis, time-lapse measurements were collected with SlideBook (3i) and analyzed in Matlab using a custom-made software (‘DotQuant’) which identifies, tracks and quantifies the fluorescent dots in each cell [28]. Statistical analysis of replication rate results was performed using Monte Carlo resampling with 1,000,000 iterations. Mitosis events were identified manually as the timepoint when the dots split and move to the mother and daughter cell.

### Chromatin immunoprecipitation (ChIP) assay

ChIP assays were performed using anti-H3K56Ac antibodies as previously described [19]. Exponentially growing cells were synchronized with 5 μg/ml α-factor for 3 hr at 25°C and then released into fresh media containing 0.2 M HU for 45 min at 25°C. Samples were collected to perform ChIP assay using H3K56Ac and H3 (as a normalization control) antibodies. The ChIP DNA was analyzed by quantitative PCR using primers (*ARS607/ARS607+14kb* and *ARS305/ARS305+12kb)* amplifying replication origins as well as fragments downstream of replication origins. The mean and SD of three independent biological replicates were shown. The percentage of ChIP DNA relative to the total input DNA was calculated (**Fig. 4**), and the ratio of H3K56Ac ChIP signal to the total H3ChIP signal was also calculated (**Fig. S2**).

### RPA foci analysis

Strains for RFA1 foci analysis were generated on the background of a BY4741 *Saccharomyces cerevisiae* strain which carries N-terminus GFP-tagged RFA1 expressed under control of its native promoter [43], as described above. To locate the cell nuclei, NLS-tetR-tdTomato was transformed into the ADE1 locus. For imaging, yeast cultures were grown overnight at 30°C to OD_600nm_= 0.2-0.6 in SC medium containing 4% glucose. Cells were fixed on microscopy slide chambers (Ibidi) coated with 2mg/ml concanavalin A (Sigma-Aldrich) and imaged using confocal microscopy as described above for 1-3h with 1 min intervals at 28°C. GFP-Rfa1 and NLS-tetR-tdTomato were excited with 488 nm and 561 nm lasers, respectively. At least 60 cells from three independent experiments were analysed for each strain and scored for the presence of one or more RPA foci during S-phase. The percentage of cells with RPA foci was estimated by dividing the number of cells exhibiting one or more RPA foci in the nucleus by the total amount of cells that were taken into account in each experiment. Results were compared with one-way ANOVA and post-hoc analysis (Dunnett’s test), after transforming the data with arcsin(sqrt(x)).

### *Can1* spontaneous mutation rate analysis

The determination of the mutation rate of the *can1* gene was performed as previously described [60] based on the Lea and Coulson method [61] with some modifications. Strains for *can1* mutation rate measurement were generated on the background of a BY4741 *Saccharomyces cerevisiae* strain as described above. Yeast cultures were grown overnight at 30°C in YPD medium to OD_600nm_ ≈ 0.8-1. Cells were then diluted 10^5^ fold, plated as single cells on SC plates and grown for 4 days at 30°C. At least 16 single colonies were randomly chosen from each strain, resuspended in 1 ml of sterile ddH_2_O and diluted to OD_600nm_ ≈ 1.0. Next, 10^5^ fold dilutions of each resuspended colony solution were plated on SC plates to define the number of viable cells and the rest of the solutions were plated on SC-Arg + 60mg/ml canavanine to obtain the number of canavanine-resistant cells. The median number of viable cells was estimated from the different colonies (replicates) and the mutation rates, the 95% Confidence Intervals (CI) and the p values were calculated by utilising the rSalvador package (v1.7) running on the R software (v3.4.3) [62].

### Rev3 inducible degradation and spot assays

Strains for spot assays were generated on the background of a BY4741 *Saccharomyces cerevisiae* strain. An auxin-inducible degron (AID) was inserted into the C’-terminus of *rev3* as previously described [28]. Briefly, a PCR fragment containing the AID sequence, the Oryza sativa TIR1 gene and the *hphMX* resistance cassette as selection marker was generated with homologous ends to the 5’ of the *REV3* yeast gene. The presence of AID results in the degradation of the fused protein after incubation of yeast cells with 1-Napthaleneacetic acid (NAA) [63,64]. The viability of *rev3-AID* histone-chaperone mutants was tested with spot assays on agarose plates. Briefly, overnight exponentially-growing yeast cultures were diluted to OD_600nm_= 0.3 and 4μ1 of 10-fold serial dilutions were spotted on YPD plates containing 200 μg/ml HYG, HYG+0.5mM NAA, HYG+0.005% MMS, or HYG+0.5mM NAA+0.005% MMS. Cells were incubated at 30°C for 32 hours and representative pictures were captured.

## Supporting Information Legends

**Figure S1: Growth defects of *cac1*-deletion strain is suppressed by *mad2-deletion.*** Growth was analyzed by a spot assay on YPD plates using a 10-fold serial dilution of yeast cultures spotted on the plate. Growth was analyzed following 24h (left) or 48h (right) of incubation.

**Figure S2: The Cac1 PIP*(F233AF234G) or Cac1 WHD*(K564EK568E) mutations lead to reduced new H3 deposition on replicating DNA.** Results are normalized to total H3 ChIP, complementing data shown in **Figure 3** (main text) indicating that loss of H3K56Ac signal is not as a result of decreased total H3.

**Figure S3: Replication times of *cac1* mutated cells measure in the presence of 20 nM of UH.** Mutations in the Cac1 WHD an PIP domains lead to increased replication times relative to WT cells in the presence of HU. Replication times of ~30.6 Kb at the vicinity of ARS413 for WT, *cac1*-deleltion, *cac1* PIP* (PIP mutant containing the F233A and F234G mutations)- and *cac1* WHD*1 (WHD mutant containing the K564E and K568E mutations)-mutant cells. The number of cells analyzed for each strain and all median values and statistical results are shown in **Table S2.**

**Figure S4: Expression of RFA1-GFP and NLS-tetR-tdTomato for validation that RPA foci are located in the cell nucleus.** Representative WT and *asf1*-deleted cells containing RPA foci are shown.

**Figure S5: The percentage of cells with RPA foci is significantly increased in *Δrad18* cells relative to WT.** Analysis of RPA foci is described in the Materials and Methods section and in **Figure 6** of the main text.

**Figure S6: Pearson correlation between percentage of cells with RPA foci and G2/M duration.** the p value is 0.017 indicating a statistically significant correlation.

**Figure S7: Depletion of DNA polymerase ζ in *cac1*- and *rtt106*-double deleted strain leads to higher sensitivity to DNA damage.** All analysed yeast strains contain *REV3* (the catalytic subunit of DNA polymerase ζ) fused to auxin inducable degron (AID) to enable its depeletion following NAA (auxin) addition to the growth media (see material and methods for details). Growth of WT, single *cac1*- or *rtt106*-deletion strains and the double *cac1*- and *rtt106*-deletion strains on plates containing YPD or YPD supplemented with 0.005% MMS were analysed with or without NAA.

**Table S1:** Parameters obtained from live cell microscopy analysis of different histone chaperone mutant strains described in the main text.

**Table S2:** Parameters obtained from live cell microscopy analysis described in the main text of different histone chaperone mutant strains in the presence of HU.

**Table S3:** Percentage of cells undergoing mitosis at a time duration of up to 90 minutes following replication of the tetR labeled array.

